# Balanced, Orientation-Dependent Dichoptic Masking in Cortex of Visually Normal Humans Measured Using Electroencephalography (EEG)

**DOI:** 10.1101/2021.09.20.461009

**Authors:** Jerry J. Zhang, Yichen Tang, Steven C. Dakin, Luke E. Hallum

## Abstract

In the human visual system, cerebral cortex combines left- and right-eye retinal inputs, enabling single, comfortable binocular vision. In visual cortex, the signals from each eye inhibit one another (interocular suppression). While this mechanism may be disrupted by e.g. traumatic brain injury, clinical assessments of interocular suppression are subjective, qualitative, and lack reliability. EEG is a potentially useful clinical tool for objective, quantitative assessment of binocular vision. In a cohort of normal participants, we measured occipital, visual evoked potentials (VEPs) in response to dichoptically-presented vertical and/or horizontal sine-wave gratings. Response amplitudes to orthogonal gratings were greater than that of parallel gratings, which were in turn greater than that of monocular gratings. Our results indicate that interocular suppression is (normally) balanced, orientation-tuned, and that suppression per se is reduced for orthogonal gratings. This objective measure of suppression may have application in clinical settings.

## Introduction

Binocular vision is a fundamental process within the primate visual system. In normally seeing humans, signals received from the left and right eyes are combined in visual cortex [1]. This combination provides the basis for single, clear, and comfortable vision. Previous work, including psychophysical studies in humans [2] and intracortical recordings in nonhuman primates [3], has shown that activity in the visual cortex arising from dichoptic presentation (i.e., when each eye views independent stimuli simultaneously) results in a mutually inhibitory effect – i.e., the signals from each eye act to reduce one another. This inhibitory mechanism, known as interocular suppression, is abnormal in individuals who suffer traumatic brain injury or who have experienced abnormal visual stimulation during development (e.g., as is the case in amblyopia) [4], [5], [6].

A common clinical test for assessing interocular suppression is the Worth Four Light Test (W4LT) [1]. In the W4LT, patients are presented with four lights (one red, two green, one white) which are viewed dichoptically using red-green goggles. The presence of suppression is determined by the patient’s report on the number of lights and their respective colours. However traditional tests such as this have several shortcomings, including a central reliance on patients making a reliable subjective report on what they perceive. This can make the test difficult to administer in preverbal children, or other non-verbal patient populations. Further, this test does not deliver an objective, quantitative measure of suppression, making it insensitive to the degree of suppression. Although other tests of interocular suppression (a.k.a. “sensory eye dominance”) have been proposed [7] that do quantify the degree of the effect, all rely on patients making a reliable subjective report. An objective, quantitative test of interocular suppression would be tremendously useful to clinicians. Furthermore, objective, quantitative measures of interocular suppression, when combined with behavioural results and computational modelling, have potential to better reveal the neural mechanisms of both normal and abnormal binocular vision.

EEG is a non-invasive, objective measure of neural activity in visual cortex. Previous psychophysical and electrophysiological studies (see Discussion) have indicated that interocular suppression in the normal visual system is orientation-dependent when invoked using a dichoptic masking paradigm [2], [8], [9], [10], [11]. Here, in a cohort of normal participants, we recorded VEPs in response to vertical and/or horizontal sine-wave gratings presented dichoptically. To investigate the effects of dichoptic masking and orientation tuning in interocular suppression, we compared VEP responses to the various dichoptic stimulus configurations.

## Methods

### Participants

We recruited seven participants (six males, one female) with normal or corrected-to-normal vision (age range: 20 to 25 years old). All participants provided written, informed consent. Protocols were approved by the University of Auckland Human Participants Ethics Committee (#021040). Prior to EEG recordings, we determined ocular dominance using a hole-in-the-card test; five (two) participants were right-eye (left-eye) dominant. Three participants then underwent full binocular assessment at the University of Auckland Optometry Clinic. These assessments revealed no significant ocular deviation (cover test, < 15 prism diopters) and no suppression or diplopia (W4LT). Two of these three participants (P4 and P6) had normal stereoacuity (Stereo Fly, ≤ 60 arc seconds), while one (P3) had abnormal stereoacuity of 160 arc seconds.

**Table 1:** Summary of vision assessment for participants P3, P4 and P6.

| **Partic.** | **Cover Test**<br>**Unaided** | **W4LT** | **Stereoacuity,**<br>**Arc Seconds** |
| --- | --- | --- | --- |
| P3 | Near, 8Δ Exophoria<br>Distance, 4Δ Exophoria | 4 Seen,<br>No Suppression or Diplopia | 160 |
| P4 | Near, 3Δ Exophoria<br>Distance, Orthophoria | 4 Seen,<br>No Suppression or Diplopia | 25 |
| P6 | Near, Orthophoria<br>Distance, Orthophoria | 4 Seen,<br>No Suppression or Diplopia | 25 |

### Dichoptic stimulation

We presented visual stimuli dichoptically using two 24-inch, gamma-corrected liquid-crystal displays [12] (LCDs; 1920×1200 px; 60 Hz; ColorEdge CG247X; EIZO, Hakusan, Japan), one on each side of the participant, and each reflected into its eye through a 45° mirror stereoscope. Viewing distance was 53 cm.

### Experimental design

Visual stimuli were vertical and/or horizontal sine-wave gratings presented within an annular aperture (inner, outer diameter = 3, 14°) with softened edges (e.g., Fig. 1). The mean luminance of the display was 60 cd/m2. Gratings were spatial frequency (SF) = 4 cyc/°, and 100% Michelson contrast. Fixation and visual attention were controlled via a discrimination task. The task is described in detail in a companion paper [13]; in brief, participants discriminated small luminance changes applied to a 0.5° disc appearing at the center of stimuli. We presented eight stimuli, illustrated in Fig. 2. Visual stimuli were programmed using MATLAB (R2019b; Mathworks, Natick, MA, USA) and the Psychophysics Toolbox (V3.0.16) [14].

**Figure 1:**
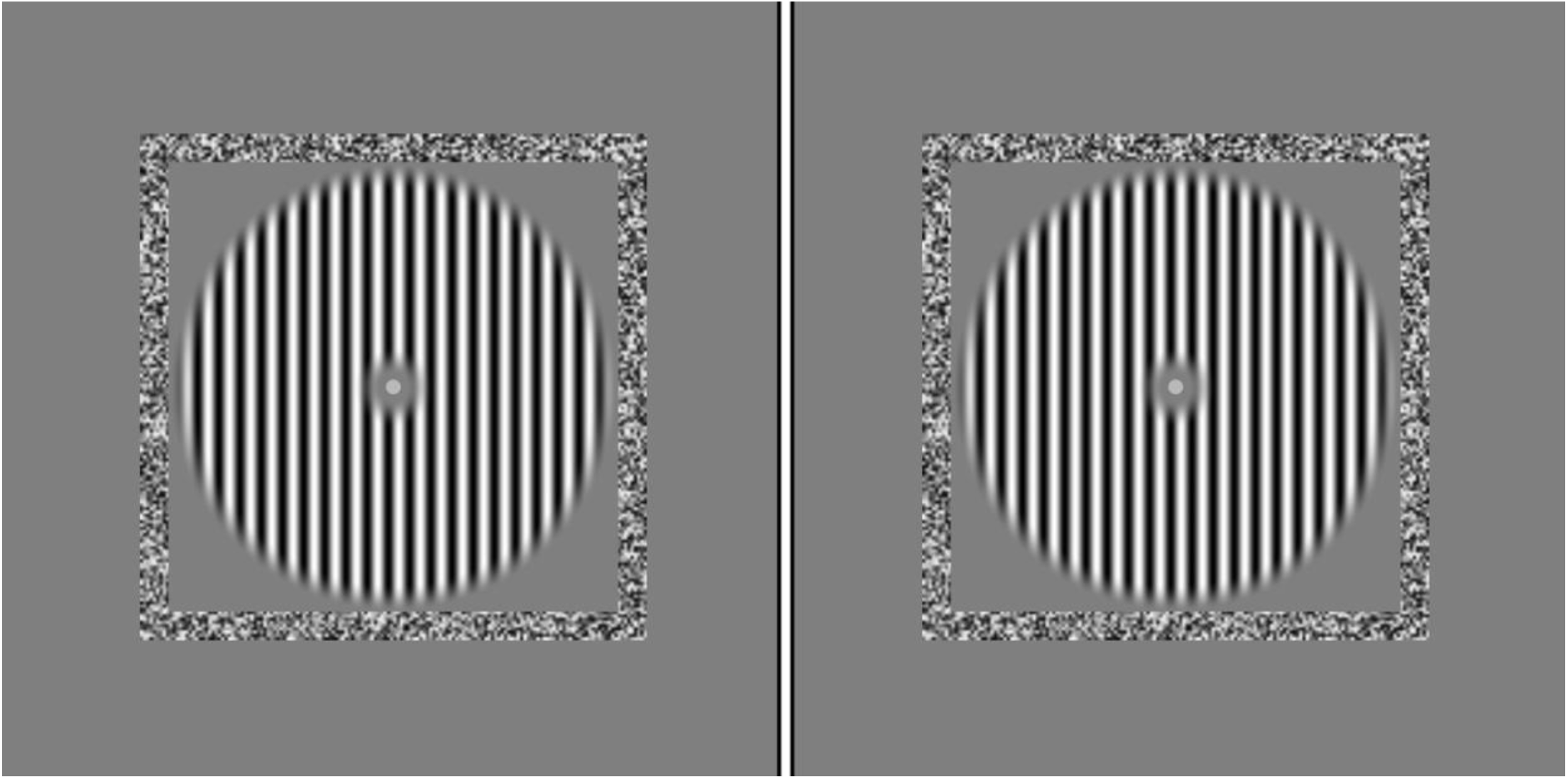
Visual stimulus. In this example the participant was presented with a “parallel” stimulus - vertical gratings to both eyes. The square white-noise border was used to enable binocular fusion. Stimuli were 4 cyc/° and the annulus outer diameter was 14°; for clarity we show low-SF stimuli.

**Figure 2:**
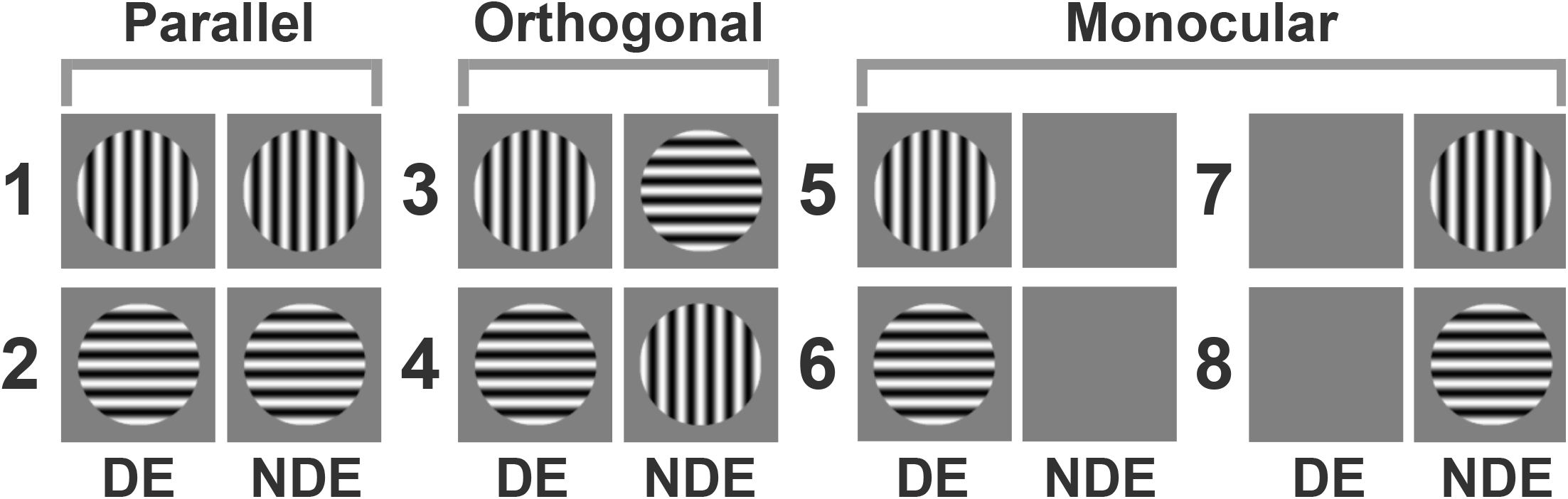
Schematic representation of the eight stimulus types used in experiments: “parallel” (types 1, 2), “orthogonal” (3, 4), and “monocular” (5, 6, 7, 8). DE, dominant eye; NDE, non-dominant eye.

Each participant sat for a total of 600 trials, recorded across five blocks (120 trials per block), with short breaks between blocks. Each block included 15 trials for each of the eight stimulus types. In each trial the stimulus was presented for 200 ms and followed by a uniform mean-luminance field for 800 ms. We pseudo-randomized the sequence of stimulation.

### EEG recording

VEPs were recorded using the ActiveTwo biopotential acquisition system (ActiveTwo MK2; BioSemi, Amsterdam, Netherlands). In collaboration with our companion study, we placed 32 electrodes in accordance with the 10-20 system [15]. The sampling frequency was 2048 Hz; signals were bandpass (2-80 Hz) and notch filtered (48-52 Hz) offline. VEPs and stimuli were aligned using a phototransistor-based trigger on the monitor.

### Data analysis

We analyzed recordings from near the primary visual cortex using occipital channel Oz, which overall produced the largest VEPs. We segmented the raw recording into 500 ms single-trial epochs, where 0 ms corresponded to stimulus onset. For each participant, we averaged epochs according to stimulus type (each average comprised 75 epochs). We extracted the absolute value of the peak amplitude in the interval from 50 to 150 ms. We used these response amplitudes to compute the following metrics: the Stimulus Dominance Index (SDI), Ocular Dominance Index (ODI) and Orientation Dominance Index (OrDI). The SDI was calculated using the following equation:

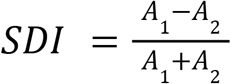

where A1 and A2 denote the response amplitudes for any two of the parallel, orthogonal or monocular stimulus types. This index varies from -1 to 1; balanced responses (i.e., where the responses to both stimuli are about equal) gives SDI = 0. OrDI and ODI were computed in the same fashion. The OrDI quantifies the balance between response amplitudes to vertical and horizontal stimuli, while the ODI quantifies the balance between response amplitudes to DE and NDE stimuli.

For statistical analysis, we used an ANOVA model comprising two fixed factors (eye, stimulus) and one random factor (participant). There were two levels to the eye factor (DE, NDE) and three levels to the stimulus factor (dichoptic, orthogonal; vertical, monocular; horizontal, monocular). In the orthogonal variable, the orthogonal stimulus for DE was defined as vertical in DE and horizontal for NDE (i.e., stimulus type 3) and the orthogonal stimulus for NDE was defined as stimulus type 4.

## Results

We recorded robust potentials from all participants. In our dataset, the responses evoked by parallel or orthogonal stimulation were greater than responses evoked by the counterpart monocular stimuli (whether NDE or DE). Furthermore, the responses evoked by orthogonal stimulation were greater than the responses evoked by parallel stimulation. This pattern of results is exemplified in Fig. 3. In the following paragraphs, we show that this finding was consistent across our cohort of seven normal participants.

**Figure 3:**
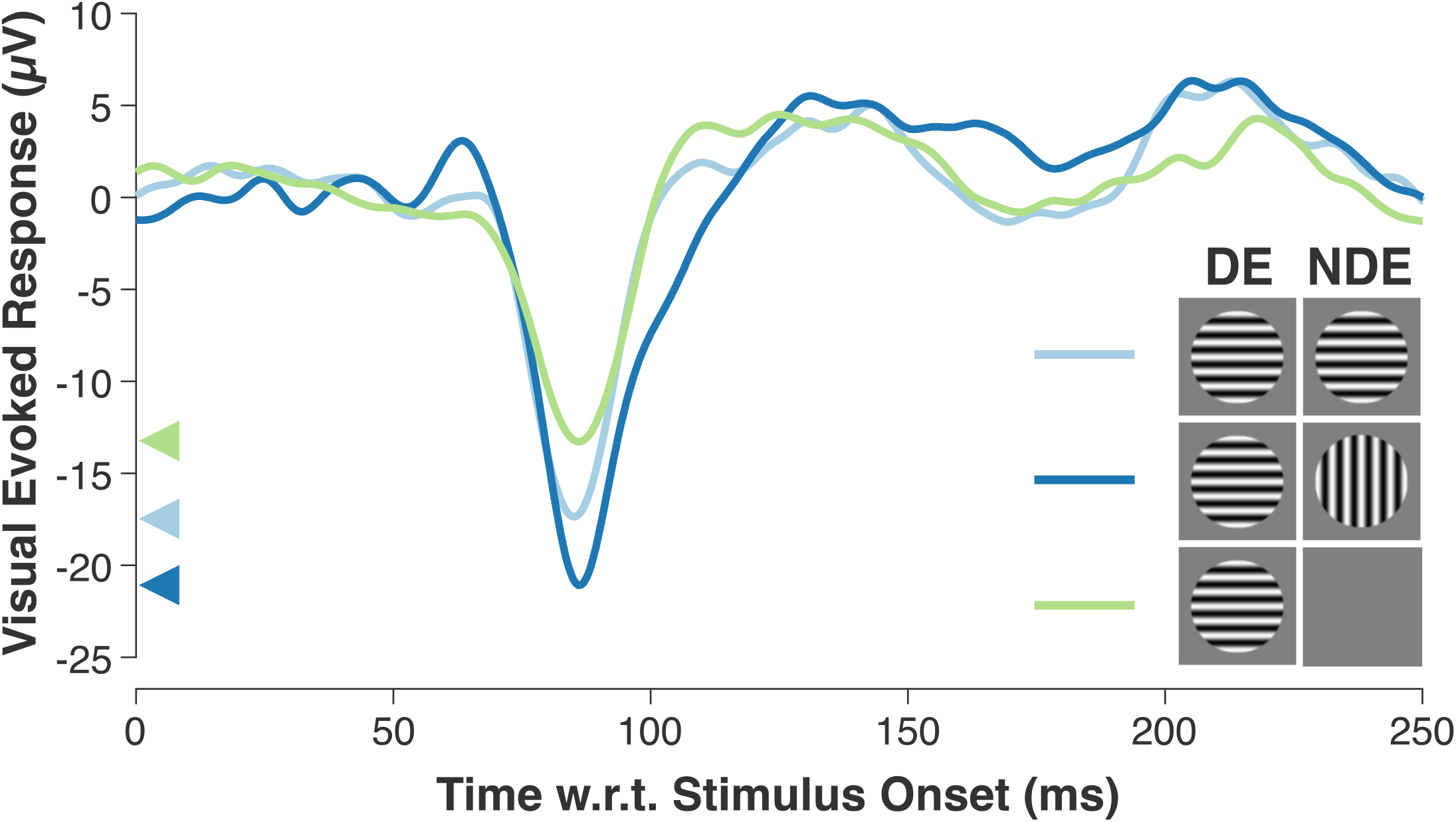
Example VEPs recorded from Participant 5. Each trace is the average of 75 trials. We show VEPs for three different stimulus types (see Fig. 2): parallel (both horizontal), orthogonal (vertical in NDE), and monocular (horizontal in DE). To quantify responses, we extracted the absolute value of peak amplitudes (arrowheads on y-axis).

### Balance in ocular dominance and orientation dominance

In all participants, responses to DE and NDE stimulation were balanced (ODI ≈ 0); this was determined by pooling over orientation (i.e., we pooled responses to stimulus type 5 with 6, and type 7 with 8) and then using a shuffle test (i.e., we shuffled the labels “DE” and “NDE”; *p* > 0.05). Similarly, in all participants, responses to vertical and horizontal gratings were balanced (OrDI ≈ 0); we pooled over DE and NDE (i.e., stimulus type 5 with 7, and type 6 with 8) and used a shuffle test (i.e., shuffling labels “vertical” and “horizontal”; *p* > 0.05).

Our ANOVA revealed that the main effect of stimulus was statistically significant (F _2,12_ = 15.94, *p* < 0.001), while the main effect of eye was not (F_1,6_ = 0.0002, *p* = 0.987). A post-hoc, paired-sample t-test of the stimulus factor showed a significant difference between orthogonal and monocular, vertical stimuli (*p* < 0.001). Likewise, there was a significant difference between orthogonal and monocular, horizontal stimuli (*p* < 0.001). There was no significant difference between vertical and horizontal monocular stimuli (*p* = 0.4765). The eye/stimulus interaction was not statistically significant (F_2,12_ = 0.1171, *p* = 0.8905).

Overall, parallel stimuli appeared to evoke larger responses than their monocular counterparts. To confirm this observation, we computed SDIs for the following comparisons (see Fig. 2): stimulus type 5 versus 1; 6 versus 2; 7 versus 1; 8 versus 2. Here, positive SDIs corresponded to larger monocular responses and negative SDIs corresponded to larger parallel responses. The first of these four comparisons – monocular (vertical in DE) versus parallel (both vertical) – revealed negative SDIs for all participants, as shown in Fig. 4a. There, error bars represent null distributions computed within participants by shuffling trial labels (“5” and “1”). The imbalance towards the parallel stimulus was statistically significant for four participants (shuffle test, *p* < 0.05). We then grouped SDIs for all four of these comparisons, as shown in Fig. 4b. These SDIs showed consistent imbalance towards parallel gratings for six of seven participants. When SDIs were pooled across participants, the mean of these 28 SDIs was -0.078, which was statistically significantly less than zero (one-sample t-test, *t* = −4.0878, *p* < 0.05).

**Figure 4:**
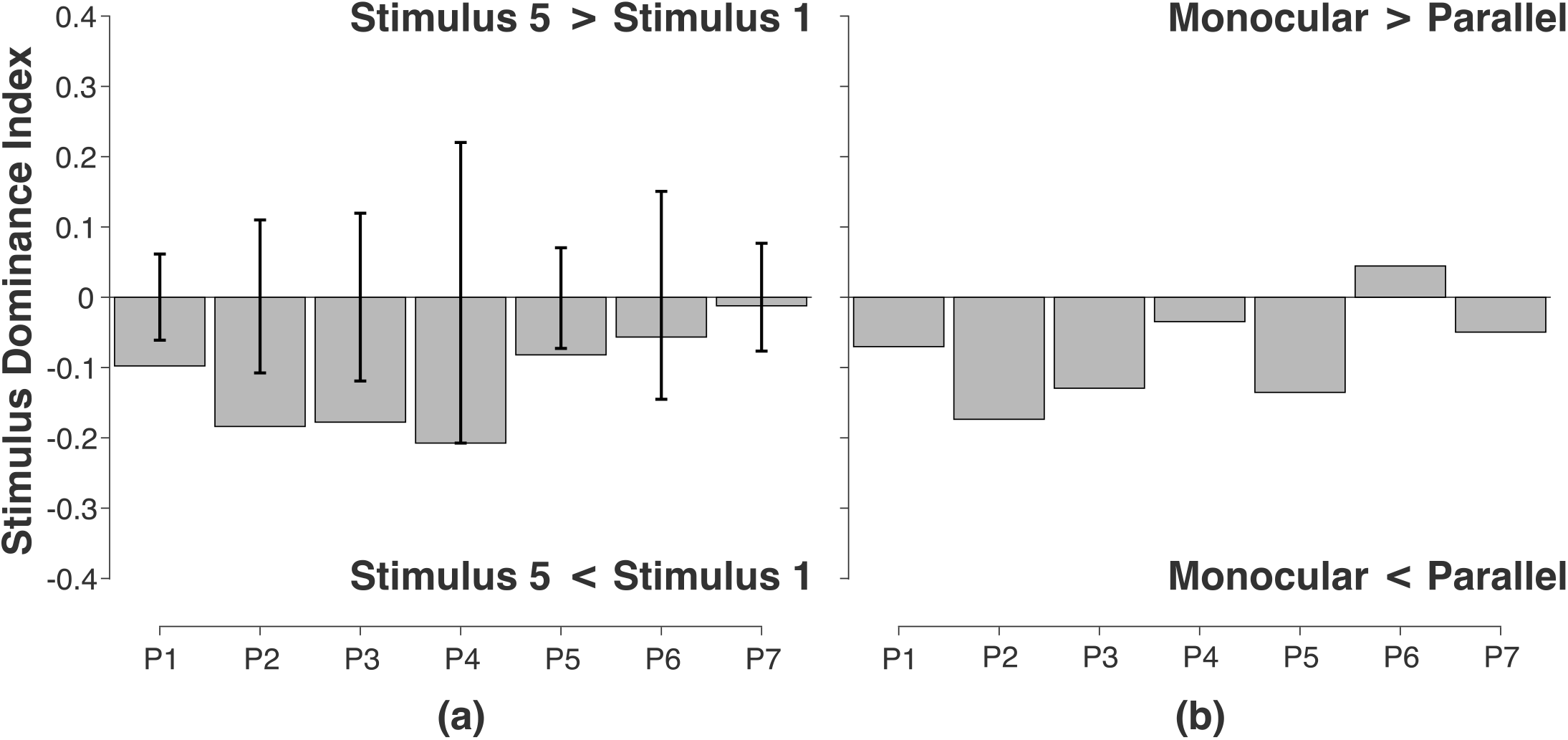
Parallel stimuli evoked larger responses than their monocular counterparts. (a) SDIs of monocular stimuli (vertical in DE) versus parallel stimulus (vertical in both eyes). Error bars represent null distributions used in a shuffle test (see text). (b) SDIs averaged within participants across four comparisons (see Fig. 2): stimulus type 5 versus 1; 6 versus 2; 7 versus 1; 8 versus 2. (P1, P2,…: Participant 1, Participant 2,…).

### Comparison of responses to orthogonal stimuli versus monocular counterparts

Overall, orthogonal stimuli appeared to evoke larger responses than their monocular counterparts. Here, we also computed SDIs for the four possible comparisons (see Fig. 2). The first of these four comparisons – monocular (vertical in DE) versus orthogonal (vertical in DE) – revealed negative SDIs for all participants, as shown in Fig. 5a. The imbalance towards the orthogonal stimulus was statistically significant for all participants (shuffle test, *p* < 0.05). We then grouped SDIs for all four of these comparisons and they showed consistent imbalance towards orthogonal gratings for six of seven participants (see Fig. 5b). When SDIs were pooled across participants, the mean of these 28 SDIs was -0.163, which was statistically significantly less than zero (one-sample t-test, *t* = −6.9517, *p* < 0.05).

**Figure 5:**
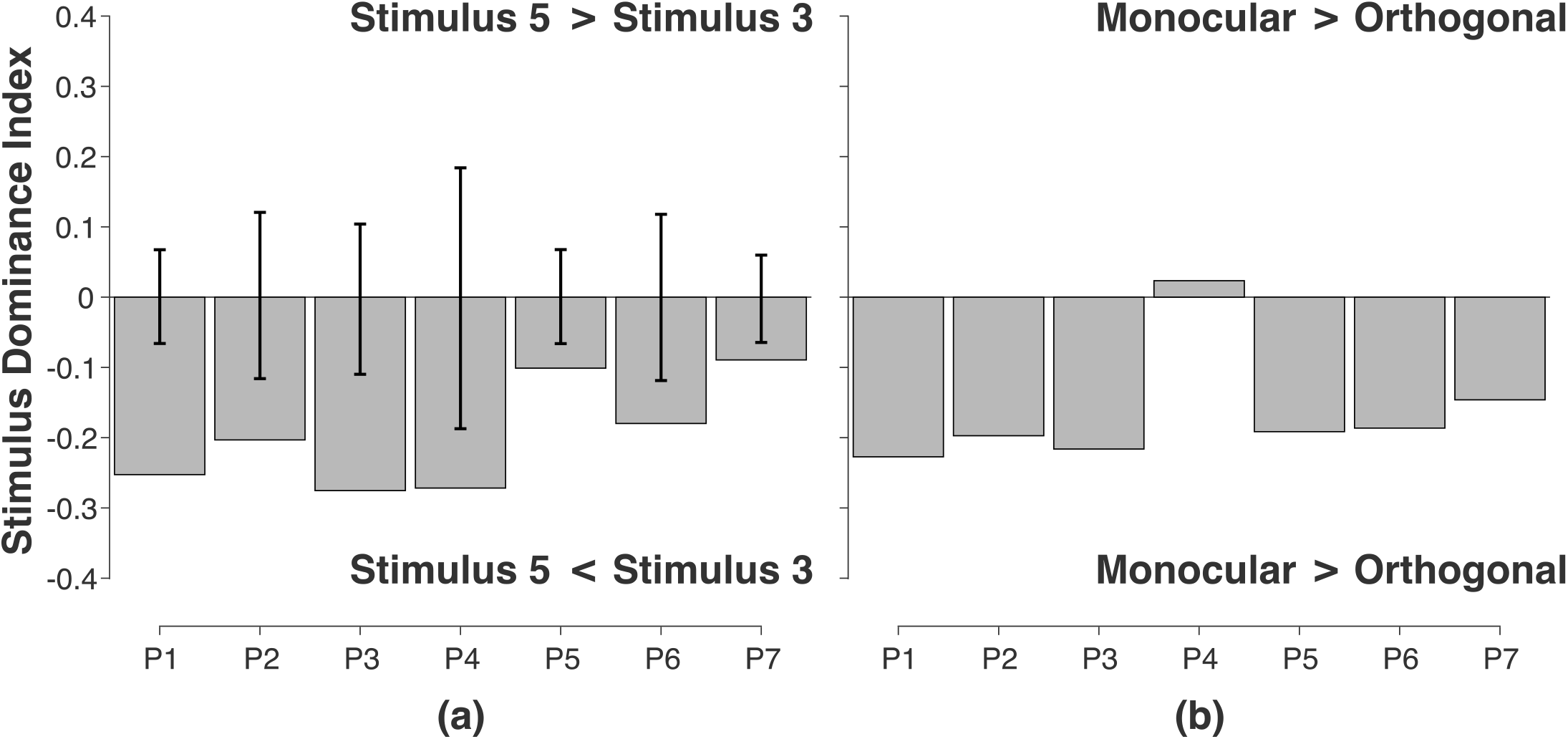
Orthogonal stimuli evoked larger responses than their monocular counterparts. (a) SDIs of monocular stimuli (vertical in DE) versus orthogonal stimulus (vertical in DE). Error bars represent null distributions used in a shuffle test (see text). (b) SDIs averaged within participants across four comparisons (see Fig. 2): stimulus type 5 versus 3; 6 versus 4; 7 versus 4; 8 versus 3.

### Comparison of responses to orthogonal versus parallel stimuli

When we compared responses to orthogonal stimuli to that of parallel stimuli, the imbalance towards the orthogonal stimulus was statistically significant for three participants (shuffle test, *p* < 0.05). We also grouped SDIs for all four of these comparisons and they showed consistent imbalance towards orthogonal gratings for six of seven participants. When SDIs were pooled across participants, the mean of these 28 SDIs was -0.086, which was statistically significantly less than zero (one-sample t-test, *t* = −3.7929, *p* < 0.05).

## Discussion

We measured VEPs in normal visual cortex. Response amplitudes to orthogonal gratings were greater than that of parallel gratings, which were in turn greater than that of monocular gratings. Our results indicated that interocular suppression is orientation-tuned, and that suppression per se is reduced for orthogonal gratings. In normal cortex, suppression appears to be balanced; DE signals suppress NDE signals, and vice versa, in roughly equal amounts.

Gong et al. [2] recently reported psychophysical findings consistent with our electrophysiological findings. They measured normal observers’ contrast detection thresholds for Gabor targets in one eye and oriented noise masks, or a uniform, mean-luminance field, in the fellow eye. Target thresholds were lowest (i.e., greatest sensitivity) for targets paired with a uniform mean-luminance field. Target thresholds were lower for orthogonal masks as compared to parallel masks, indicating a release from suppression for orthogonal stimuli. When target and mask were swapped between the eyes, threshold elevations were similar, indicating balance. Taken together, their results and ours both indicate interocular suppression is orientation-dependent and balanced.

Three previous studies [9], [10], [11] measured VEP responses to dichoptic stimuli in normal participants, but the results of these studies were not in agreement with each other. Harter et al. [9] measured responses to line patterns, orientation = θ, in the RE while the LE was continuously stimulated with a 0° (vertical) line pattern. As θ increased, so too did the VEP response amplitude, indicating a release from interocular suppression. Jakobsson et al. [10] measured responses to contrast-reversing sine-wave gratings, calculating the ratio of dichoptic-to-monocular VEP amplitudes. That ratio was greater when gratings were orthogonal as compared to parallel. Our results confirm and extend Harter’s and Jakobsson’s: we controlled and diverted visual attention, neither eye was in an adapted state and our design enabled the assessment of ocular and orientation balance. Tyler and Apkarian [11], however, found conflicting results. In a single participant, their dichoptic, orthogonal response amplitudes were less than that of the monocular counterpart. Roeber [16] recently measured, at occipital and parietal sites lateral to Oz, dichoptic VEP responses. In future work, we aim to compare our lateral measurements to his.

In summary, we found balanced, orientation-dependent interocular suppression in normal cortex. In future work, we aim to measure interocular suppression in a cohort afflicted with injury or disease, and further consider EEG’s utility for clinical binocular assessment.

## Acknowledgements

We thank Emanuele Romano and Kathryn Sands for technical and clinical support, respectively.

## Notes

### Competing Interest Statement

The authors have declared no competing interest.

